# Testing the Efficacy of Transungual Drug Delivery System (Nail Lacquer) Containing Terbinafine Against *Aspergillus brasiliensis* for the Topical Treatment of Onychomycosis

**DOI:** 10.1101/2022.02.03.478980

**Authors:** Shivani. H. Rao, Mangesh. R. Bhalekar, Samruddhi. H. Kelkar, Shivani. S. Godbole, Riddhesh. A. Kharche

## Abstract

According to a survey of 42 epidemiological studies, the cases of Non-dermatophyte mould onychomycosis (NDMO) caused by Aspergillus species have been found to increase. The number ranges from <1 to 35% of all cases of onychomycosis in the general population and the number is higher in the diabetic population. Around7.7-100% of cases of NDMO are caused by Aspergillus species. Hence the attempt of investigation to test the efficacy of the drug by transungual drug permeation has been made. The antifungal agent ‘Terbinafine’ was incorporated in a topical dosage form (Nail lacquer). The formulation is optimized by a drug design matrix of 2^3^ type using Design-Expert®software, version 11 (DX11). The antifungal assay was performed using the Kirby Bauer disk diffusion method and other evaluation parameters like Drying time, Viscosity are evaluated. Optimized formulation with the highest desirability was selected out of all the possible formulations which showed the best results with good physicochemical properties and antifungal activity. Thus, it can be concluded that medicated nail lacquers containing Terbinafine can be used as a successful method to treat non-dermatophyte mould onychomycosis (NDMO) caused by Aspergillus species.

## Introduction

Onychomycosis also called Tinea unguium is a fungal infection of the nails that causes discoloration, thickening, and separation from the nail bed. [1] Onychomycosis occurring in younger population is less while its occurrence in an older population above 60 years is Maximum. It is caused by various organisms like *Aspergillus, Trichophyton, Epidermophyton*, Yeast, *Fusarium* & *Candida* species. Among these *Aspergillus* are increasingly being reported as the causative agent. [2] [3] *Aspergillus* species not being keratinophilic like other Dermatophytes cause onychomycosis as a secondary infection after an external injury. *Aspergillus* species infects the proximal part of the nail and fungus invading proximally accumulates the debris along with nail turning white distally.[4]

*Aspergillus brasiliensis* is one of the members of black *Aspergillus* organism group which was earlier recognised as *Aspergillus niger* and renamed as *Aspergillus brasiliensis* in 2010. *A. brasiliensis* being thermotolerant indicates that it can literally be grown everywhere even at extreme conditions. [5] *A. brasiliensis* can cause otomycosis and severe onychomycosis in already immune deficient or healthy people. It is responsible for nail discolouration into black and milky white base colour along with periungual inflammations. On microscopic examination of the nail, dichotomous septate hyphae formation was observed. [6] [7]

Terbinafine is an orally and topically active antifungal agent which belongs to the chemical class of Allylamines. Terbinafine is highly lipophilic hence it is widely distributed in tissues and has a long elimination half-life.[8] [9] Oral administration leads to first pass metabolism of the drug and in such cases patients are advised to have liver function tests performed to reduce risk of liver injury whereas in case of a novel drug delivery system (NDDS) such as nail lacquer, first pass metabolism is avoided and bioavailability is increased. [10] [11] Terbinafine inhibits the biosynthesis of ergosterol by blocking the action of squalene epoxidase which results in ergosterol depleted fungal cell membrane and toxic accumulation of intracellular squalene leading to fungicidal activity. [12] [13] The minimum inhibitory concentration (MIC) of terbinafine against *A. brasiliensis* was found to be 0.4 μg/ml.[14]

## Nail Lacquer

It has been observed that maximal antifungal efficacy against *Aspergillus brasiliensis* is achieved by formulating medicated nail lacquers. [15] [16] The nail lacquer forms a film on the nail surface which acts as a depot that leads to optimized, sustained diffusion and allows continuous penetration of the Active Pharmaceutical Ingredient (API) to achieve high tissue concentration which is required to treat onychomycosis. [17]

The following study was conducted to formulate and evaluate Terbinafine hydrochloride nail lacquer for the treatment of onychomycosis.

## Materials and Method

The nail lacquer was formulated using: Nitrocellulose (RS grade with viscosity range 0.25 to 0.5 cps), Toluene, Dibutyl phthalate, Ethyl alcohol, Benzyl alcohol, Polyethylene glycol (PEG)-400 (all chemicals used were of analytical grade.) The fungal strain used to test the efficacy of Terbinafine was *Aspergillus brasiliensis* (National Collection of Industrial Micro-organisms (NCIM) Pune; ATCC 16404, GenBank Accession: KR908766).

### Preparation of fungal suspension

*A. brasiliensis* was sub-cultured using Potato Dextrose Agar (PDA) (Media: Difco™) under aseptic conditions and Potato Dextrose Broth (PDB) (Media: Difco™) was used to prepare fungal suspension for evaluating zone of inhibition by the Kirby Bauer method.[18]

### Formulation of nail lacquer

Simple mixing

The nail lacquer was prepared by simple solution method. The nitrocellulose (film former) was dissolved in dibutyl phthalate (plasticizer) and was mixed the solvent system comprising of ethyl alcohol and benzyl alcohol, to this toluene (co-solvent) was added since it aids in dissolution of plasticizer and film former followed by addition of PEG-400 (viscosity enhancer). Lastly Terbinafine (API) was incorporated by sonication. [19]

The nail lacquer formula was optimized by employing 2^3^ full factorial design. (Table 1). A total of eight formulations were developed along with a negative control. Concentration of nitrocellulose, solvent system and PEG-400 were selected as independent variables. Zone of inhibition, drying time and viscosity were taken as dependent variables. Data obtained were then subjected to Design-Expert^®^ software, version 11 (DX11), (Stat-Ease, Inc., Minneapolis, USA) and analysed statistically. 3D response plot was generated to estimate simultaneous influence of nitrocellulose, solvent system (Ethyl alcohol: Benzyl alcohol) and PEG-400 on dependent variables.

**Table 1:** Factor combinations and levels for 2^3^ Factorial design

| Level | Nitrocellulose (g) | Solvent System |  | PEG-400 (ml) |
| --- | --- | --- | --- | --- |
|  |  | Ethyl alcohol (ml) | Benzyl alcohol (ml) |  |
| +1 | 4 | 12.32 | 6.16 | 1.4 |
| -1 | 2 | 9.24 | 3.08 | 0.6 |

**Table 2:** Composition of trial runs for optimization of nail lacquer

| Formulation number | Nitrocellulose (A)<br>(g) | Solvent system (B)<br>(ml) |  | PEG-400 (C)<br>(ml) | Toluene (ml) | Dibutyl phthalate (ml) | Terbinafine (g) |
| --- | --- | --- | --- | --- | --- | --- | --- |
|  |  | Ethyl alcohol | Benzyl alcohol |  |  |  |  |
| F1 | 2 | 9.24 | 6.16 | 0.6 | 2.6 | 2 | 0.4 |
| F2 | 4 | 9.24 | 6.16 | 0.6 | 2.6 | 2 | 0.4 |
| F3 | 2 | 12.32 | 3.08 | 0.6 | 2.6 | 2 | 0.4 |
| F4 | 4 | 12.32 | 3.08 | 0.6 | 2.6 | 2 | 0.4 |
| F5 | 2 | 9.24 | 6.16 | 1.4 | 2.6 | 2 | 0.4 |
| F6 | 4 | 9.24 | 6.16 | 1.4 | 2.6 | 2 | 0.4 |
| F7 | 2 | 12.32 | 3.08 | 1.4 | 2.6 | 2 | 0.4 |
| F8 | 4 | 12.32 | 3.08 | 1.4 | 2.6 | 2 | 0.4 |
| Control<sup>1</sup> | 4 | 9.24 | 6.16 | 1.4 | 2.6 | 2 | 0.4 |

<sup>a</sup>Control- The control (negative control) is used to assess the preparation batch for possible contamination during the preparation and processing steps. The control formulation is processed along with and under the same conditions as the associated samples to include all steps of the analytical procedure.

## In vitro antifungal activity

Disk diffusion antifungal sensitivity test (Kirby Bauer method). [20] [21]

Punched out disks of Whatman^®^ ashless filter paper (Grade no-42), pore size-2.5 μm; were made and sterilized using moist heat sterilization (autoclaving) at 121°C and 15 psi for 15 minutes. The filter paper punches are then impregnated with 5 μl of prepared nail lacquer and was left to dry for 24 hours. The fungal suspension was prepared using PDB (dissolve 0.6 g in 25 ml purified water and heat to dissolve the powder thoroughly and sterilized by using moist heat sterilization technique at 121°C and 15 psi for 15 minutes). The fungal strain was inoculated in 20 ml PDB under aseptic conditions and placed in a rotary orbital shaker for 24hr at 37°C and 90 RPM. The disk diffusion method was carried out using PDA (Dissolve 0.78 g in 20 ml distilled water and heated gently to avoid sedimentation of starch and sterilized by autoclaving at 121°C and 15 psi for 15 minutes), poured 25 ml in a petri plate and allowed to solidify. The aforementioned procedure was repeated for 5 petri plates. Furthermore, 10 μl of prepared fungal suspension were measured and inoculated by spread plate technique using a glass spreader in the petri plates. The prepared Whatman filter paper punches were placed inverted (the face containing formulation downwards) on the media using sterilized forceps. The procedure was repeated for all the formulations followed by incubation for 24 hours at 37°C.

## Evaluation of nail lacquer

Evaluation of the developed nail lacquer was carried out as per Bureau of Indian Standards, IS9245:1994 (10).

## Determination of drying time

A film of sample was applied with the help of nail brush on a clean glass slide. Time taken for film to dry (dry-to-touch) was noted down using a stopwatch. (Dry-to-touch is a condition at which the film may be touched with a clean fingertip without the resultant transfer of any material to the finger.) [22]

## Determination of Viscosity

Accurately weighed 20 g of the sample was placed in beaker and viscosity of optimized formulation measured by Brookfield viscometer at 60 rpm 25 °C. [23] [24]

## Zone of inhibition

The measurement of the zone of inhibition was done by physical methods such as using a metric ruler. The zone was identified by visual inspection and the radius and the diameter of the zone was measured by placing the scale above the petri dish and the value was read manually. Larger the diameter of the zone, greater is the effectiveness of the antifungal agent. [25]

## Statistical analysis and validation of experimental design

Effect of independent variables on dependent variables was optimized by Design-Expert^®^ Software version 11 (DX11). One way ANOVA was applied and polynomial equation was generated for zone of inhibition, drying time and viscosity. Validation of experimental design was done using polynomial equation. The predicted value was calculated from reduced polynomial equation and compared with experimental value.[26] [27]

## Result

Attempts for testing the efficacy of the Transungual Drug Delivery System (Nail Lacquer) containing Terbinafine against *Aspergillus brasiliensis* for the topical treatment of Onychomycosis were made and the following parameters were studied-

## Statistical analysis and validation of experimental design

Effect of independent variables nitrocellulose, solvent system (Ethyl alcohol: Benzyl alcohol) and PEG-400 on dependent variables: zone of inhibition, drying time and viscosity were assessed statistically using Design-Expert^®^ software version 11 (DX11). The mathematical equations generated for the response parameter(s) are expressed as:

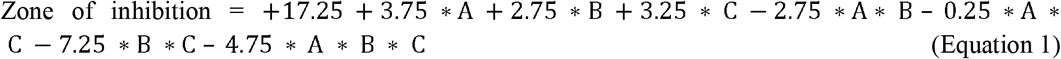

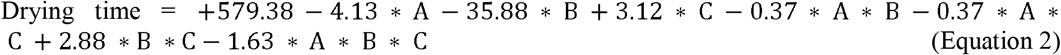

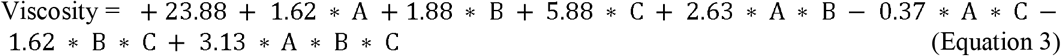

Considering these polynomial equations, the responses can be predicted at any value of Nitrocellulose, solvent system (Ethyl alcohol: Benzyl alcohol) and PEG-400. Zone of inhibition values obtained were between 2 to 36 mm. The study was performed according to CLSI guidelines and following observations were obtained (Table 3) which were compared with the standards provided by CLSI (Table 4). According to the standards, the formulations; F6, F4 and F7 showed zones of inhibitions of 36mm, 30mm and 20mm respectively, proving their effectiveness against the fungal strain. The formulation F3 showed considerable effectiveness by showing the zone of inhibition to be 18mm (Fig.4). The remaining formulations viz. F1, F2, F5, F8 did not pass the test thus proving to be ineffective. [28] [29] However all the factors had effect on developing zone of inhibition. The values increased on simultaneously increasing the concentration of nitrocellulose solvent composition ad also the PEG. Similar results were obtained earlier. [30]

**Table 3:** Experimental data for zone of inhibition of nail lacquer formulations (F1-F8 and Control)

| Formulation | Zone of Inhibition (mm) | Rank | Comment |
| --- | --- | --- | --- |
| F1 | 2 | 7 | Resistant |
| F2 | 6 | 7 | Resistant |
| F3 | 18 | 4 | Intermediate |
| F4 | 30 | 2 | Susceptible |
| F5 | 14 | 5 | Resistant |
| F6 | 36 | 1 | Susceptible |
| F7 | 20 | 3 | Susceptible |
| F8 | 12 | 6 | Resistant |
| Control | 0 | 7 | - |

**Table 4:** Interpretive category of zone of inhibition according to CLSI guidelines.

| Interpretive category | Zone diameter (mm) |
| --- | --- |
| Susceptible | $\geq 20$ |
| Susceptible-Dose dependent | 15-19 |
| Intermediate | 15-19 |
| Resistant | $\leq 14$ |

**Table 5:** Experimental data for drying time and viscosity of nail lacquer formulations (F1-F8 and Control)

| Formulation | Drying time (s) | Viscosity (cps) |
| --- | --- | --- |
| F1 | 620 | 12 |
| F2 | 610 | 17 |
| F3 | 540 | 20 |
| F4 | 535 | 23 |
| F5 | 618 | 34 |
| F6 | 613 | 25 |
| F7 | 556 | 23 |
| F8 | 543 | 37 |
| Control | 518 | 20 |

The increase in concentration of polymer and high boiling solvent helped formation of a good non-sticky film which leads to better drug release and permeation (Equation 1). However, at high nitrocellulose concentration the zone of inhibition increased steeply with ethanol concentration (Fig.1). Similarly, drying time decreased with the increasing concentration of ethyl alcohol (i.e., solvent system) and polymer concentration (Equation 2), However, the PEG-400 being liquid and non-volatile caused an increase in drying time (Fig.2). The viscosity was positively affected by all the three independent variables (Equation 3). The desired viscosity was obtained in range on decreasing the concentration of PEG-400 (Fig.3).

**Fig.1-.**
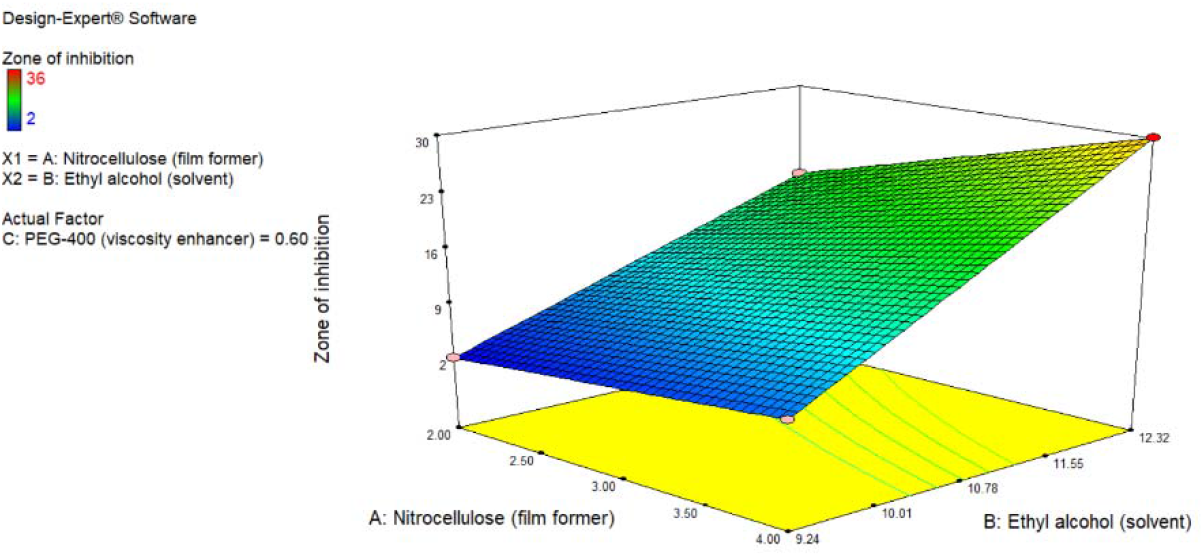
3D response surface plots showing the effect of independent variable-zone of inhibition.

**Fig.2-.**
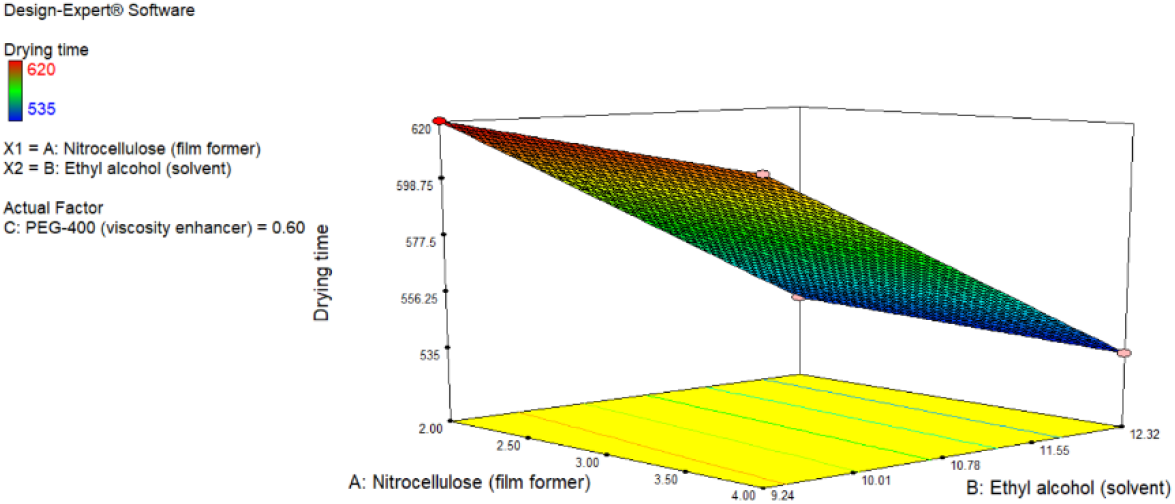
3D response surface plots showing the effect of independent variable-drying time.

**Fig.3-.**
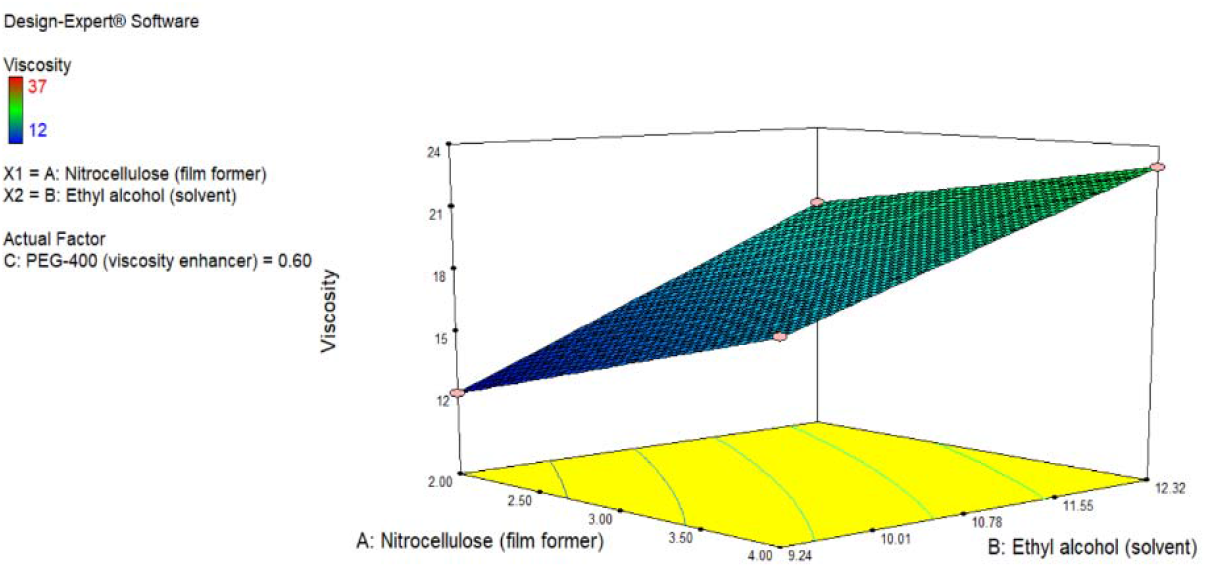
3D response surface plots showing the effect of independent variable-viscosity.

**Fig.4-.**
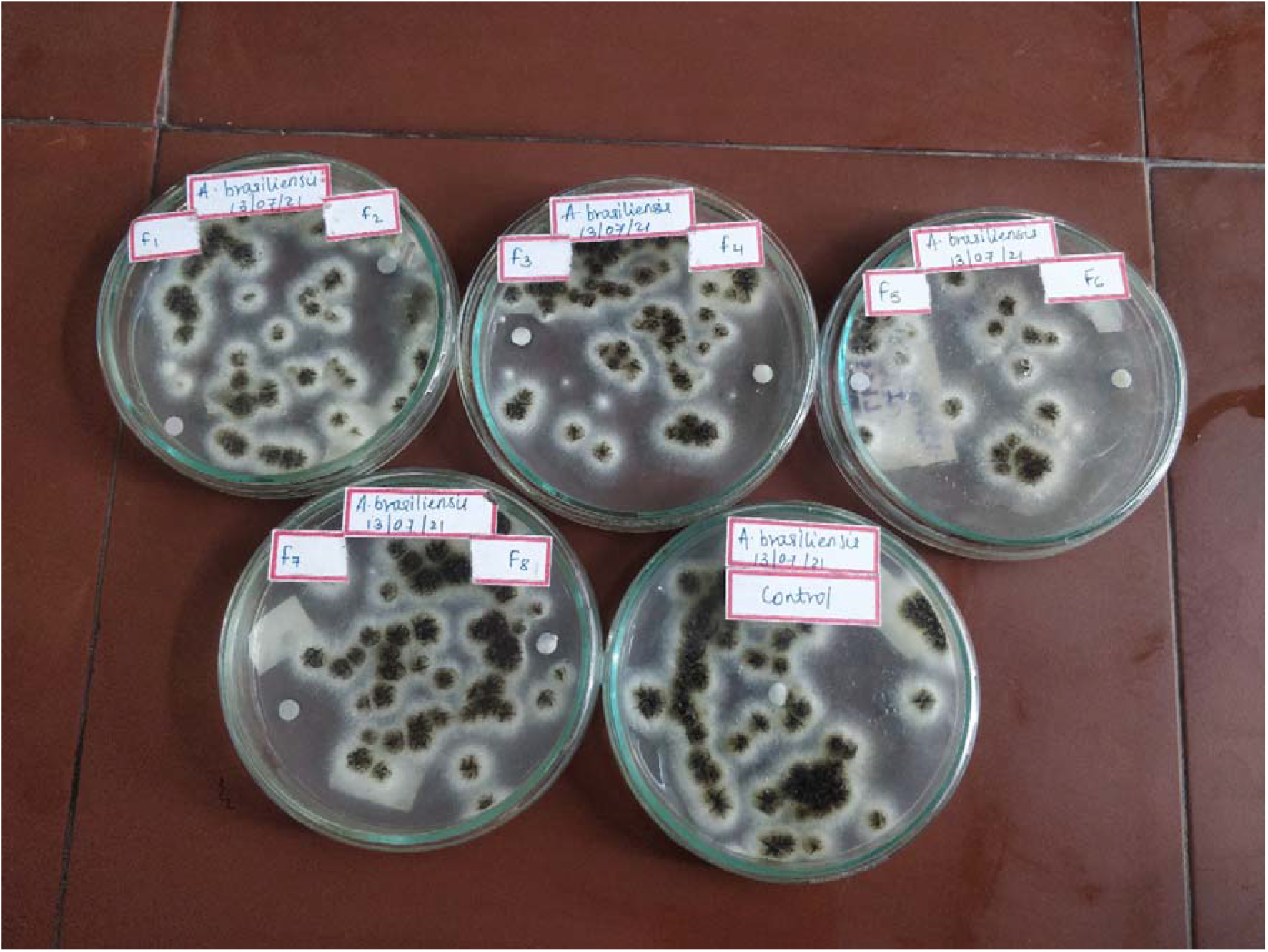
in-vitro antifungal activity of nail lacquer formulation (F1-F8) and control.

Finally, the optimized formulation was selected on the basis of desirability. From the polynomial equation the predicted values of zone of inhibition, drying time and viscosity were calculated as 29.9999 mm, 536 s and 23.4 cps respectively. Experimental values were found to be 30 mm, 535 s and 23 cps respectively. The desirability for the optimized formulation was presented to be 0.907.

## Discussion

The systemic therapy with antifungal drug (Terbinafine) shows non-optimal results hence efforts are made to incorporate Terbinafine as a topical agent in the form of nail lacquer against NDMO caused by *A. brasiliensis*. The nail lacquer was tested for its antifungal efficacy by Kirby Bauer method-the filter paper was used to simulate the nail on which the nail lacquer was applied, the antifungal agent diffused through the filter paper onto the inoculated media to elicit its effect. Other parameters such as drying time and viscosity were also studied. The result obtained indicated that the formulation F4 was identified to be optimum on the basis of maximum desirability (0.907). (F4 containing 4 g nitrocellulose, solvent system in the ratio of 4:1 (Ethyl alcohol: Benzyl alcohol), 0.6 ml PEG-400 along with 2% Terbinafine). Thus, it can be concluded that medicated nail lacquers containing Terbinafine can be used as a successful method to treat non-dermatophyte mould onychomycosis (NDMO) caused by *Aspergillus* species.

## Acknowledgements

The authors would like to thank AISSMS College of Pharmacy, Principal and management. The authors are thankful to Emcure Pharmaceuticals Ltd. Pune, for providing Terbinafine Hydrochloride as a gift sample and National Collection of Industrial Micro-organisms (NCIM) Pune, for supplying the fungal strain of *A. brasiliensis*.

## Statements and Declarations

### Author Contributions

All authors contributed to the study conception and design. Material preparation, data collection and analysis were performed by Mrs. Shivani Rao, Dr. Mangesh Bhalekar, Miss Samruddhi Kelkar, Miss Shivani Godbole and Mr. Riddhesh Kharche. The first draft of the manuscript was written by Miss Samruddhi Kelkar, Miss Shivani Godbole and all authors commented on previous versions of the manuscript. All authors read and approved the final manuscript.

## Ethical Statement

The authors declare that there were no animal studies or clinical trials involving human subjects performed as a part of the experiment, thus this research is exempted from ethics approval.

